# The Macroevolution of Filamentation Morphology Across the *Saccharomycotina* Yeast Subphylum

**DOI:** 10.1101/2025.11.26.690719

**Authors:** Christina M. Chavez, Marie-Claire Harrison, Thodoris Danis, Marizeth Groenewald, Chris Todd Hittinger, Antonis Rokas

**Affiliations:** Department of Biological Sciences, Vanderbilt University, Nashville, TN 37235, USA; Evolutionary Studies Initiative, Vanderbilt University, Nashville, TN 37235, USA; Westerdijk Fungal Biodiversity Institute, 3584 Utrecht, The Netherlands; Laboratory of Genetics, DOE Great Lakes Bioenergy Research Center, Wisconsin Energy Institute, Center for Genomic Science Innovation, J.F. Crow Institute for the Study of Evolution, University of Wisconsin-Madison, WI 53726, USA

**Keywords:** evolutionary cell biology, hyphae, pseudohyphae, *Saccharomycotina*, ancestral state reconstruction, machine learning, convergent evolution

## Abstract

*Saccharomycotina* yeasts are a highly diverse and widely distributed subphylum of ascomycete fungi that exhibit diversity in their asexual growth morphologies; the human commensal yeast *Candida albicans* exhibits both unicellular and filamentous growth, whereas the baker’s yeast *Saccharomyces cerevisiae* grows unicellularly by budding. Filamentous growth (and the production of hyphae) can comprise linear and branched budding cells that do not undergo cell separation, termed pseudohyphae, or of tubular filaments with septa that perforate allowing movement of organelles, termed true hyphae. We integrated phenotypic, genomic, metabolic, and isolation environment data from 1,154 strains from 1,051 *Saccharomycotina* species to examine the variation and evolutionary history of filamentation across *Saccharomycotina* and determine the extent to which genes, metabolic traits, and environments can predict filamentation types. We found that 63.37% of strains can form filaments; 6.56% can form true hyphae, 42.40% can form pseudohyphae, and 14.39% can form both true hyphae and pseudohyphae. We found that the distributions of species that can produce true hyphae or that can filament were more strongly correlated with the yeast phylogeny than the distribution of species with pseudohyphae. Ancestral state reconstruction suggested that the ability to form true hyphae and/or pseudohyphae evolved several times in yeast evolution. We infer that most yeast ancestors likely produced pseudohyphae or lacked filaments, while the *Saccharomycotina* last common ancestor likely produced pseudohyphae but not true hyphae. Machine learning models trained on genomic and metabolic features predicted filament morphologies with about 70% accuracy. Connecting the evolution of *Saccharomycotina* morphologies to their genomic, metabolic, and ecological characteristics will enrich our understanding of the evolution of the diversity of lifestyles observed in this lineage.

## Introduction

Fungi are a diverse group of eukaryotic organisms that exhibit unicellular (yeast) and/or multicellular (filamentous) morphologies during vegetative growth (Powers-Fletcher et al., 2016) In general, filamentous growth is important for response to environmental changes and stress, aids with biofilm formation, and increases nutrient acquisition. For example, filamentous fungi in soil environments may transfer water through their filamentous mycelial networks (Pérez-Izquierdo et al., 2021). Induction of filamentous growth can be especially advantageous to organisms in nutrient-poor and variable environments (Berman, 2006; Karasz et al., 2022). For example, several pathogenic fungi can thrive in host environments by utilizing filamentous morphologies during stressful conditions, such as changes in pH, temperature, and nutrients (Chow et al., 2021; Du et al., 2020).

*Saccharomycotina* is an ancient subphylum of fungi that contains approximately 1,200 known species (Groenewald et al., 2023; Opulente et al., 2024). Even though *Saccharomycotina* are commonly referred to as yeasts, several species in the subphylum can also exhibit filamentous growth (Chavez et al., 2024; Kurtzman et al., 2011). Filamentous growth in *Saccharomycotina* manifests itself through two different types of filaments or hyphae, termed pseudohyphae and true hyphae. Pseudohyphae are chains of budding cells in which the daughter cells have stayed attached to their mother cells. True hyphae are highly polarized tubular cells with septae that allow for perforation of organelles or multinucleation. In *Saccharomycotina* that can filament, one can typically observe both yeast cells and filaments (pseudohyphae and/or true hyphae) in the same culture. In some species, the switch between the different types of growth can depend on the environment. For example, environmental stresses, such as changes in temperatures and low glucose availability, are associated with the production of filaments in human hosts for pathogenic species, such as *C. albicans* and *Candidozyma auris* (synonym *Candida auris*) (Chow et al., 2021; Du et al., 2020). Furthermore, in certain culture media, *C. albicans* yeast cells form at 25°C, pseudohyphae at 30°C, and true hyphae at 37°C (Mukaremera et al., 2017). Finally, some *Saccharomycotina* species always or nearly always exhibit filamentous growth (and seldom or never grow as yeasts), including several species in genera across the order *Dipodascales*, such as *Dipodascus, Geotrichum,* and *Magnusiomyces* (Kurtzman et al., 2011).

The genetic pathways required for filamentation have been extensively studied in species such as *S. cerevisiae* and *C. albicans* (Cullen & Sprague, 2012), and they are related to the cell polarity and biofilm formation pathways (Bi & Park, 2012; Chiou et al., 2017). An important genetic pathway that allows for reorganization of cell polarity to produce filaments is the MAPK signaling pathway. There are many genes that are required for both the cell polarity and MAPK signaling pathways; for example, regulators of the polarisome bind active GTPase Cdc42p to establish a site of polarity for cell growth, and regulator kinases recruit Cdc42p to activate downstream MAPK signaling. Further, a MAPK kinase, Fus3p, phosphorylates Bni1p, a component of the polarisome; when the *FUS3* gene is deleted cell polarization does not function properly (Matheos et al., 2004). Cell polarity allows for the growth of budding cells, as well as pseudohyphae and true hyphae. When filamentous cells grow continuously, they can form biofilms (Váchová & Palková, 2018). Biofilm formation increases cell-cell adhesion using cell surface proteins, such as Flo11p, which is one of the flocculins required for filamentation in *S. cerevisiae* (Van Mulders et al., 2009; Verstrepen & Klis, 2006). Expression of *FLO11* is also under the control of the MAPK and Snf1 pathways (Laxman & Tu, 2011; Van Mulders et al., 2009; Verstrepen & Klis, 2006).

To examine how a filamentous lifestyle has evolved across *Saccharomycotina*, we cataloged the variation and distribution of filament types (pseudohyphae and true hyphae) across the subphylum, examined their phylogenetic conservation, and inferred their evolutionary history. We also employed machine learning to identify genomic, metabolic, and/or environmental traits that are associated with the observed variation in filamentation across *Saccharomycotina* yeasts.

## Results

### Inter- and intra-ordinal variation of filamentation across *Saccharomycotina* yeasts

To elucidate the variation of filament morphology, we collected all available filament descriptions from reference works on yeast taxonomy (Kurtzman et al., 2011; theyeasts.org) and calculated the frequency of filamentation-associated traits across the 12 taxonomic orders in the *Saccharomycotina* phylogeny. Of the 1,154 yeast strains, 133 (11.53%) lack information about filamentation. From the remaining 1,021 yeasts, 374 / 1,021 (36.63%) are known to grow only unicellularly, whereas 647 / 1,021 (63.37%) can generate filaments. Among these strains, 433 of 1,021 (42.41%) can only form pseudohyphae, 67 / 1,021 (6.56%) can only form true hyphae, while 147 / 1,021 (14.40%) are polymorphic and can form both pseudohyphae and true hyphae (Figure 1). Interestingly, 435 / 1,021 strains (3.33%) are known to only grow via filamentation, i.e., they are not known to exhibit yeast growth or form buds but instead may generate conidia, blastoconidia, or arthroconidia. Most species of the genera *Dipodascus*, *Geotrichum, Galactomyces,* and *Magnusiomyces* (all from the order *Dipodascales*) produce true hyphae that disarticulate into arthroconidial cells. Three *Blastobotrys* species (order *Dipodascales*) generate primary and secondary conidia from true hyphae. Other species that form blastoconidia and / or arthroconidia were found in the orders *Dipodascales* (genus *Saprochaete*) and *Ascoideales* (*Ascoidea rubescens*)*. Dipodascus geniculatus* (order *Dipodascales)* and *Dipodascopsis tothii (*order *Lipomycetales*) do not produce yeast cells nor arthroconidia, and *Eremothecium cymbalariae* (order *Saccharomycetales*) is described as “purely hyphal” (Kurtzman et al., 2011).

**Figure 1.**
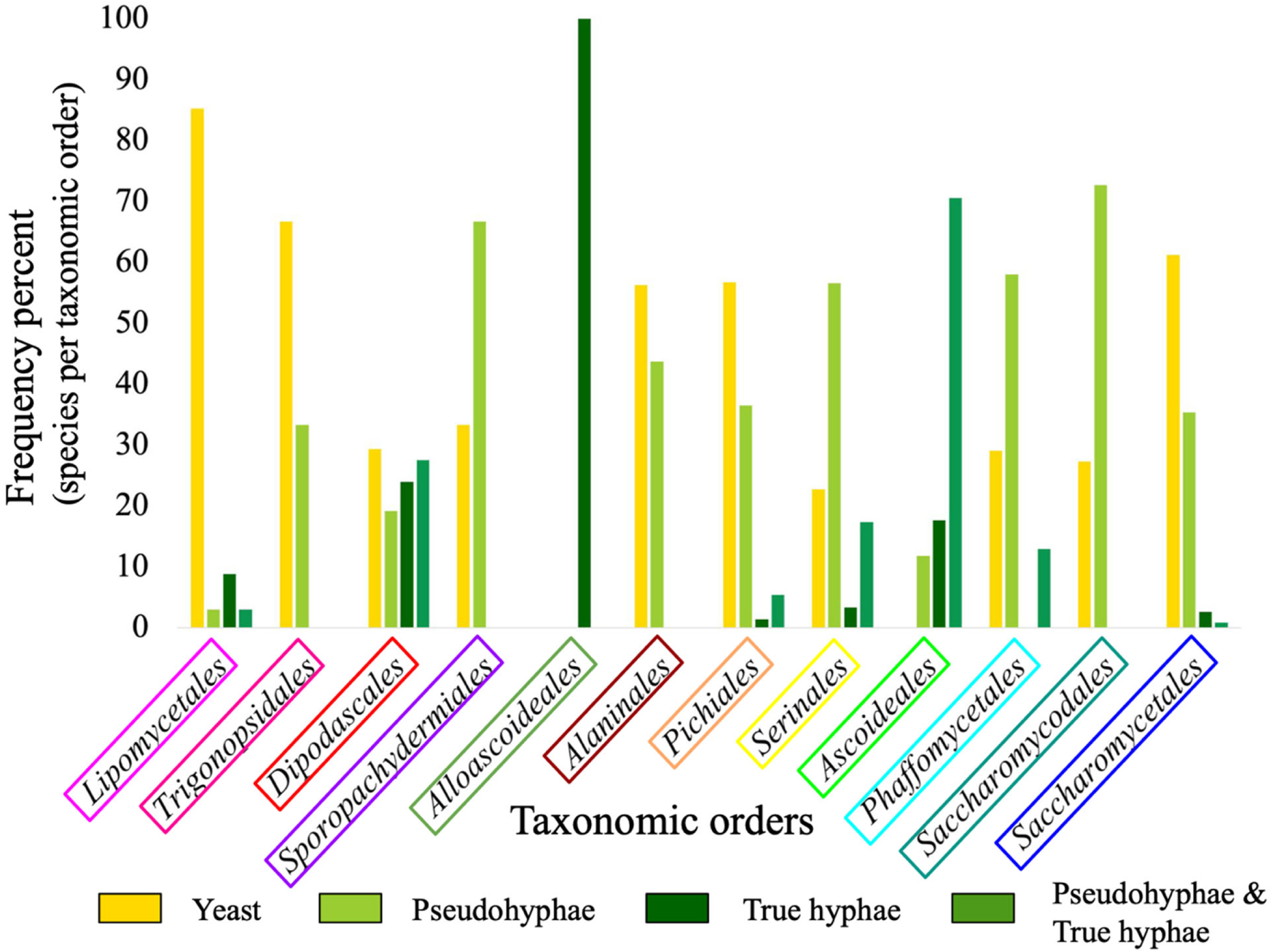
Variation in the frequency of filamentation types across the 12 *Saccharomycotina* orders. Histograms of the frequency of filamentation trait types, including filaments absent (i.e., unicellular or yeast growth; yellow), pseudohyphae present (light green), true hyphae present (dark green), and polymorphic pseudohyphae and true hyphae (medium green). To obtain the frequency values for each order, the count of each filamentation type was divided by the total number of species in the order for which data are available. The numbers of yeasts analyzed in each order were: *Lipomycetales*: 33, *Trigonopsidales*: 15, *Dipodascales*: 174, *Sporopachydermiales*: 3, *Alloascoideales*: 3, *Alaninales*: 14, *Pichiales*: 145, *Serinales*: 384, *Ascoideales*: 17, *Phaffomycetales*: 94, *Saccharomycodales*: 22, and *Saccharomycetales*: 118.

The *Saccharomycotina* orders with the highest frequency of filamentation are *Alloascoideales* (100%; 3 / 3 tested strains in the order), *Ascoideales* (80.95%; 17 / 21), *Serinales* (69.53%; 299 / 430), *Saccharomycodales* and *Sporopachydermiales* (66.67%; 16 / 24 and 2 /3, respectively). *Lipomycetales* has the lowest frequency of filamentation (14.71%; 5 / 34), followed by *Trigonopsidales* and *Saccharomycetales* (33.33%; 5 / 15, and 63.38%; 45 / 71, respectively).

Almost all taxonomic orders of *Saccharomycotina* contain strains that produce pseudohyphae, apart from *Alloascoideales*. Orders *Saccharomycodales* (72.73%; 16 / 22 tested strains in the order), *Sporopachydermiales* (66.67%; 2 / 3), *Serinales* (56.59%; 219 / 387), and *Phaffomycetales* (58.06%; 54 / 93) have the highest frequencies of yeasts that can form pseudohyphae. The orders with the lowest frequencies are *Dipodascales* (19.16%; 32 / 167), *Ascoideales* (11.76%; 2 /17), and *Lipomycetales* (2.94%; 1 / 34).

Only 67 / 1,021 tested strains in the order (6.56%) of *Saccharomycotina* yeasts representing seven of the 12 orders are known to produce true hyphae. These are *Dipodascales* (23.95%; 40 / 167), *Ascoideales* (17.65%; 3 / 17), *Lipomycetales* (9.09%; 3 / 33), *Serinales* (3.39%; 13 / 384), *Saccharomycetales* (2.59%;3 / 116), *Pichiales* (1.36%; 2 / 148), and all three *Alloascoideales* yeasts.

Yeasts from seven different orders are known to generate both true hyphae and pseudohyphae. These are (from highest to lowest) *Ascoideales* (70.59% tested strains in the order; 12 / 17), *Dipodascales* (27.55%; 46 / 167), *Serinales* (17.97%; 69 / 384), *Phaffomycetales* (12.90%; 12 / 93), *Pichiales* (5.41%; 8 / 148), *Lipomycetales* (2.94%; 1 / 34), and *Saccharomycetales* (0.86%; 1 / 116). *Dipodascales* is arguably the order with the highest levels of interspecific variation of filamentation, (23.95%; 40 / 167 yeasts) produce true hyphae, (19.16%; 32 / 167) produce pseudohyphae (27.55%; 46 / 167) produce both pseudohyphae and true hyphae, and (29.34%; 49 / 167) are not known to generate filaments. A few other orders exhibit substantial interspecific variation, including *Serinales*: more than half generate pseudohyphae (56.59%; 219 / 387 yeasts), others generate both pseudohyphae and true hyphae (17.31%; 67 / 387), a small percentage generates true hyphae (3.36%; 13 / 387), and more than 20% are not known to form filaments (22.74%; 88 / 387). In contrast, other orders exhibit lower levels of interspecific variation; for example, most *Saccharomycodales* yeasts are known to form pseudohyphae (72.73%; 16 / 22) and the remaining ones do not produce filaments (27.27%; 6 / 22). Thus overall, there is diversity of filament types both within, but especially between orders with respect to types of filamentation.

### Filamentation morphologies tend to be evolutionarily conserved

To determine if the pattern of variation in filamentation reflects the *Saccharomycotina* phylogeny, we calculated the phylogenetic signal D (Fritz & Purvis, 2010) of each morphological trait. Values of D near 1 reflect phylogenetic randomness (i.e., trait distribution is random with respect to phylogeny), whereas values of D near 0 reflect phylogenetic conservation (i.e., closely related species exhibit more similar traits than species that are more distantly related). The presence/absence of filamentation (including presence/absence of only true hyphae, of only pseudohyphae, and of both true hyphae and pseudohyphae) hyphae produced by a strain had a high phylogenetic signal (D value = 0.26, probability of D value resulting from Brownian phylogenetic structure p-value = 0.002), the presence/absence of both pseudohyphae and true had a slightly lower phylogenetic signal (D value = 0.33, p-value = 0.003), whereas the presence/absence of pseudohyphae had an even lower phylogenetic signal (D-value = 0.53, p-value = 0). The distribution of the presence/absence of true hyphae, despite its low D-value, was not significant (D-value = 0.02, p-value = 0.456), likely due to the relatively small number of yeasts can generate true hyphae. These results suggest that filamentation-related traits are generally, but not always, phylogenetically conserved and that closely related species tend to exhibit similar filamentation morphologies.

### Evolutionary history of *Saccharomycotina* reveals filamentous ancestors

To estimate the evolutionary history of the different filamentation traits, we performed ancestral state reconstruction analyses using different models of trait evolution (see Methods). We analyzed our filamentation data as a single character with three-character states: YEAST, PSEUDOHYPHAE, and TRUE HYPHAE. We found that the model with the best fit (i.e., lowest AIC value) was the ARD model, which assumes that each transition rate is independent of one another and can have different values in the gain and loss of a trait state (Figure 2A). We found that the evolutionary transition rate from yeast to pseudohyphae [(YEAST → YEAST + PSEUDOHYPHAE = 0.007) + (YEAST + PSEUDOHYPHAE → PSEUDOHYPHAE = 0) = 0.007] was slightly higher than the transition rate from yeast to true hyphae [(YEAST → YEAST + TRUE HYPHAE = 0) + (YEAST + TRUE HYPHAE → TRUE HYPHAE = 0.003) = 0.003]. Similarly, the transition rate from pseudohyphae to yeast [(PSEUDOHYPHAE → PSEUDOHYPHAE + YEAST = 0) + PSEUDOHYPHAE + YEAST TO YEAST = 0.008) = 0.008] was higher than the transition rate from true hyphae to yeast [(TRUE HYPHAE → TRUE HYPHAE + YEAST = 0.002) + (TRUE HYPHAE + YEAST → YEAST = 0) = 0.002]. Furthermore, transition rates from filamentous to yeast morphologies were typically higher than those from yeast to filamentous morphologies, consistent with the macroevolutionary transition of the *Saccharomycotina* predominant yeast lifestyle from a filamentous ancestor over evolutionary time.

**Figure 2.**
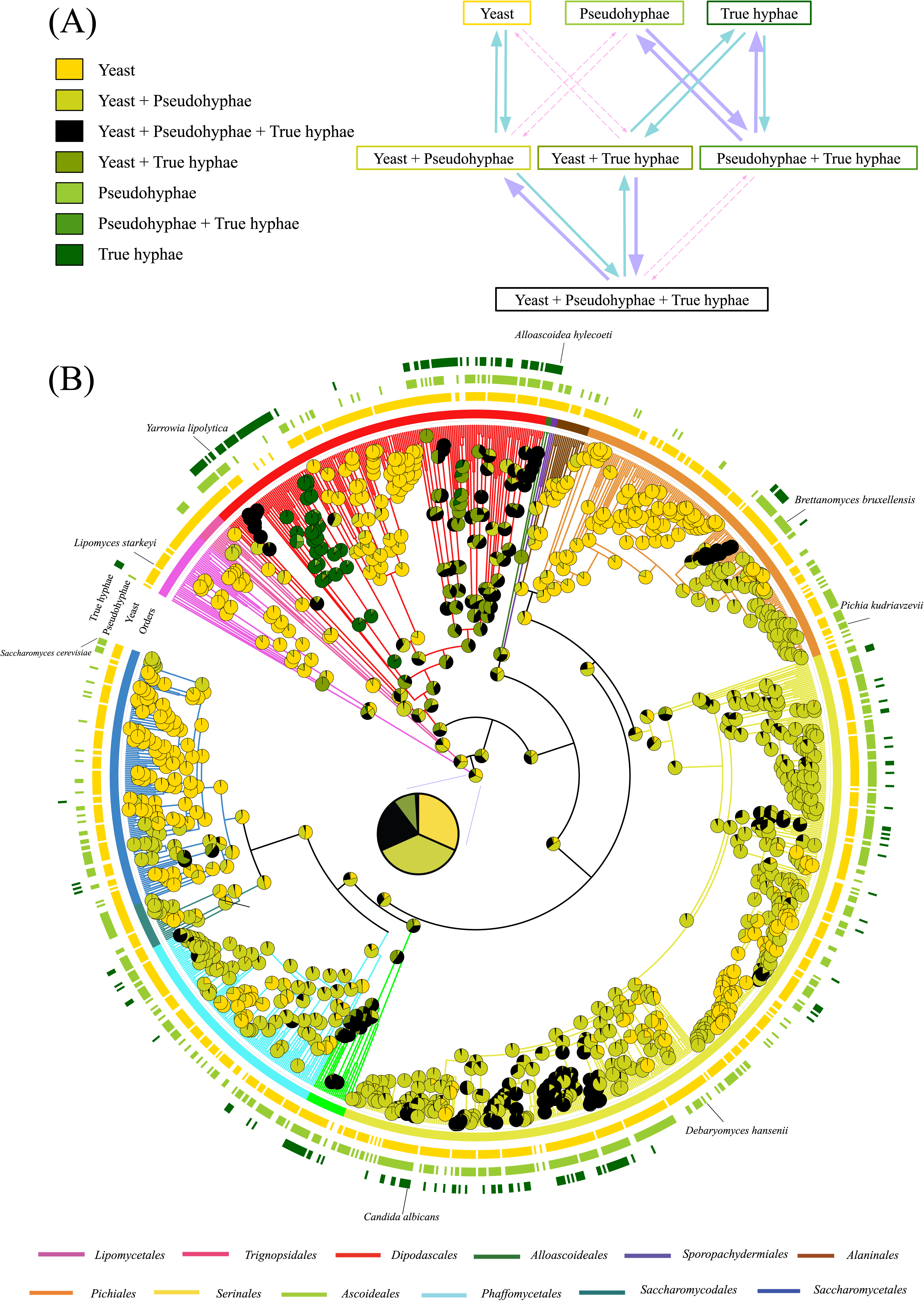
The last common ancestor of *Saccharomycotina* yeasts could likely grow filaments. Transition rates of filamentation trait states and ancestral state reconstruction of filamentation types across *Saccharomycotina* yeasts. A) The transition rate map of the filament trait states with rates as arrows; thicker arrows represent higher transition rates and thinner arrows represent lower rates. The color of arrows represents varying transition rate ranges including 0.01 - 0.099 (purple arrows), 0.001 - 0.009 (green arrows), and rates of 0 (pink dotted arrows). B) Ancestral state reconstruction of filamentation types. Pie charts in ancestral nodes correspond to the likelihood-inferred ancestral state(s). The enlarged pie chart within the phylogeny is the enlarged ancestral state inference for the *Saccharomycotina* last common ancestor. The tips of the phylogeny represent 1,021 extant yeasts that have filament trait states. The colored circles around the phylogeny show (from inner to outer) the taxonomic order, yeast (yellow), pseudohyphae (light green), and true hyphae (dark green) trait states of each yeast species. For example, a *Dipodascales* species that can produce yeast, pseudohyphal, and true hyphal morphologies is shown as red in the first inner circle (reflecting its order), yellow in the second (reflecting its ability for yeast growth), light green in the third (pseudohyphal growth), and dark green (true hyphal growth).

The ability to form filaments has evolved five times independently; two of these transitions included pseudohyphae, and the three other transitions included true hyphae. Our inference suggests that most *Saccharomycotina* ancestors produced filaments, and they seldomly could only grow as yeasts (Figure 2B). The *Saccharomycotina* last common ancestor likely had the ability to filament and was more likely to form pseudohyphae than true hyphae.

### Genomic and metabolic traits predict the presence of filamentation variation across *Saccharomycotina*

Filamentation is likely due to multiple genetic and environmental factors (Basso et al., 2019; Chow et al., 2021; Cullen & Sprague, 2000). To examine how variation in genomic, metabolic, and environmental features was associated with the variation of filament types observed across the subphylum, we used three recently generated data matrices of genomic features (which included variation in InterProScan IDs for all predicted orthologous genes, gene families, and domains) (Opulente et al., 2024), metabolic growth features (presence/absence of growth in different substrates and environments) (Harrison et al., 2024; Kurtzman et al., 2011; Opulente et al., 2018), and isolation environment features (the descriptions for isolation environments from Kurtzman et al. 2011 of each strain were converted to an environmental trait matrix using an ecological ontology) (M. Harrison et al., 2024; Opulente et al., 2024).

To examine the relationship between filamentation traits and genomic, metabolic, and environmental features, we trained a random forest machine learning algorithm on the three data matrices independently and tested how well each of these data matrices predicted filamentation. We found that genomic features had moderate accuracy (71%) when predicting filament presence / absence across 1,021 *Saccharomycotina* yeast species (Figure 3A). Accuracy was lower when the algorithm was trained on metabolic features (67%; Figure 3B), and slightly above random (57%) when trained on environmental isolation features (Figure 3C). The low accuracy achieved with environmental isolation features is expected as the isolation environment may be different than the environment that most commonly contains that strain.

**Figure 3.**
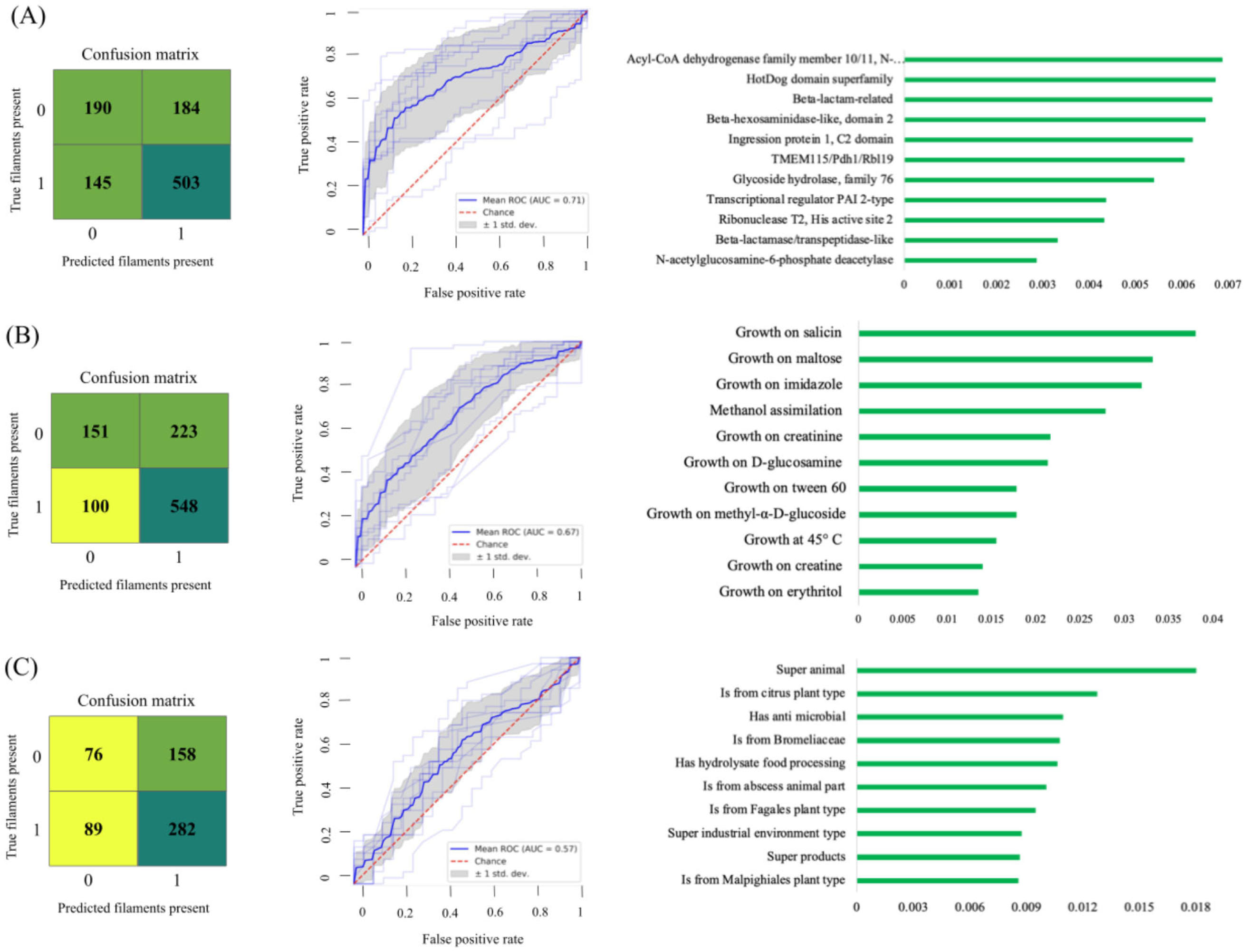
The top genomic, metabolic, or environmental features that predict filament morphology across *Saccharomycotina* yeasts. A.) The best machine learning model was trained on genomic data (InterPro ortholog presence / absence); its accuracy, measured by the receiver operating characteristic (ROC) area under the curve (AUC), was 0.71. The top ten InterPro orthologs that best predicted filaments had a range of importance from 0.0029 to 0.0069. B.) The trained on metabolic data (presence / absence of ability to grow on different substrates) had an accuracy value of 0.67. The top ten metabolic traits that best predicted filaments had a range of importance from 0.014 to 0.038. C.) The model trained on isolation environment data had an only slightly higher than random accuracy of 0.57. The top ten isolation environment traits that best predicted filaments had a range of importance from 0.018 to 0.0086.

Top genomic features that predicted filament production best according to Gini impurity included InterPro features of N-acetylglucosamine metabolism, fatty acid metabolism, and cell polarity. The top two InterPro features were Acyl-CoA dehydrogenase (InterPro ID IPR041726) and HotDog domain of thioesterases and dehydratases (InterPro ID IPR029069), both involved in fatty acid metabolism (Figure 3A). For *S. cerevisiae* and *C. albicans*, fatty acid metabolism, specifically beta-oxidation, is important for growth in low nutrient environments (Wang et al. 2024; Otzen et al. 2014). Gene features related to *N*-acetylgucosamine metabolism, including *N*-acetylglucosamine-6-phosphate deacetylase (InterPro ID IPR003764) and glucosamine-6-phosphate (InterPro ID IPR004547) (*C. albicans* gene *NAG1*), were also among the top InterPro features. *N*-acetylgucosamine has been implicated as a signal during formation of hyphae in *S. cerevisiae* (order *Saccharomycetales*)*, C. albicans* (order *Serinales*), and *Yarrowia lipolytica* (order *Dipodascales*) ( Basso et al., 2019; Du et al., 2020; Pérez-Campo & Domínguez, 2001). Other interesting top InterPro features were the CDC24 domain (InterPro ID IPR033511) and the Ras-GTPase activating domain (InterPro ID IPR001936). A daughter cell can begin growth once establishment of polarity occurs which involves the GEF Cdc24p, and other members of the cell polarity network that bind at the plasma membrane to regulate of the accumulation of Cdc42p (Chiou et al., 2017; Diepeveen et al., 2017).

Given the importance of cell polarity in filamentation and that the process has been very well studied in *S. cerevisiae* (Bi & Park, 2012; Diepeveen et al., 2017), we further examined how cell polarity- and filamentation-related genes predicted the absence and presence of filamentation. We found 20 InterPro features in the top 500 genomic features. In addition to CDC24 and GTPase activating proteins (GAPs), InterPro features related to cell polarity included actin patch protein, and cyclins. An InterPro feature required for filamentation included the flocculin Flo11 domain (InterPro ID IPR018789) (Cullen & Sprague, 2000; Lambrechts et al., 1996), which is present in Flo11p, an important protein for biofilm formation (Van Mulders et al., 2009).

The top 10 metabolic features that best predicted filamentation included growth on different carbon sources (e.g., erythritol, methanol, maltose) and nitrogen sources (e.g., creatine and imidazole) (Figure 3B). Limitation of carbon and nitrogen sources can induce filamentous growth in *S. cerevisiae* and *C. albicans* (Cullen & Sprague, 2012; Du et al., 2020). Three α-glucosides (i.e., salicin, maltose, and methyl-α-D-glucoside) were among the top 10 features. Interestingly, recent studies have shown that the ability to grow on numerous α-glucosides is significantly associated with the size of the gene family encoding α-glucoside transporters (AGTs), a gene family with a rich history of gene duplications across *Saccharomycotina* (Crandall et al., 2024; David et al., 2025). Notably, several α-glucosides have been shown to induce filamentation in *S. cerevisiae* (Van De Velde & Thevelein, 2008). Other top metabolic features that were predictors of filamentation included growth in succinate, which has been previously found to promote growth in *Magnusiomyces magnusii* (order *Dipodascales*) that is known to only generate septate hyphae (Il’chenko et al., 2005), and amyloid, or starch, formation, which occurs during the production of filamentous biofilms in species such as *C. albicans* (Mourer et al., 2021). Finally, the top isolation environment features associated with filamentation were “animals” and “isolation from *Citrus* plants” (Figure 3C).

## Discussion

Our study describes the variation of filamentation, and the genomic and metabolic features associated with it across the *Saccharomycotina* subphylum. Even though *Saccharomycotina* organisms are commonly referred to as “yeasts”, we found that most species in the clade (63.37%) can filament. However, filamentation ability varies considerably across *Saccharomycotina*, especially between taxonomic orders (Figures 1 and 2). Both the ability for filamentous growth and for pseudohyphal growth were phylogenetically conserved, suggesting that closely related species tend to exhibit similar cellular morphologies (Figure 2).

Past examinations of representatives from diverse fungal clades suggest that the ability to produce filaments is ancient and has been marked by repeated acquisitions of the yeast lifestyle within various lineages, such as *Saccharomycotina, Pezizomycotina,* and *Taphrinomycotina* (Harris, 2011; Kiss et al., 2019). In this study, through ancestral state reconstruction of the filamentation morphologies of over 1,000 species, we infer that the *Saccharomycotina* last common ancestor was most likely capable of filamentous growth (Figure 4). Interestingly, the last common ancestor most likely produced pseudohyphae (Figure 2B); taxa with the ability to generate true hyphae likely originated from ancestors capable of pseudohyphal growth (Figure 2A). Given that *Saccharomycotina* yeasts emerged from ancestors that grew as true hyphae (e.g. the Dikarya last common ancestor) and the true hyphal growth of species in the sister subphylum *Pezizomycotina*, it is perhaps surprising that the *Saccharomycotina* ancestor is inferred to have been capable of pseudohyphal growth rather than true hyphal growth. Studies of the molecular mechanisms governing filamentation in *C. albicans* have shown that true hyphal growth mechanistically differs from pseudohyphal growth in both the polarized growth mechanisms involved and in cell cycle organization (Sudbery, 2011). However, the differences in mechanisms are largely regulatory, and numerous mutations have been described that result in true hyphal growth, instead of pseudohyphal growth, and vice versa (Sudbery, 2011), suggesting that there are likely numerous mutational paths that can give rise to true hyphal growth. Furthermore, comparative genomics studies have shown that the genetic machinery governing filamentation, whether involving pseudohyphae or true hyphae, is ancient (Wu et al., 2022) and has been evolutionarily retained, albeit in reduced form, in *Saccharomycotina* yeasts (Kiss et al., 2019).

**Figure 4.**
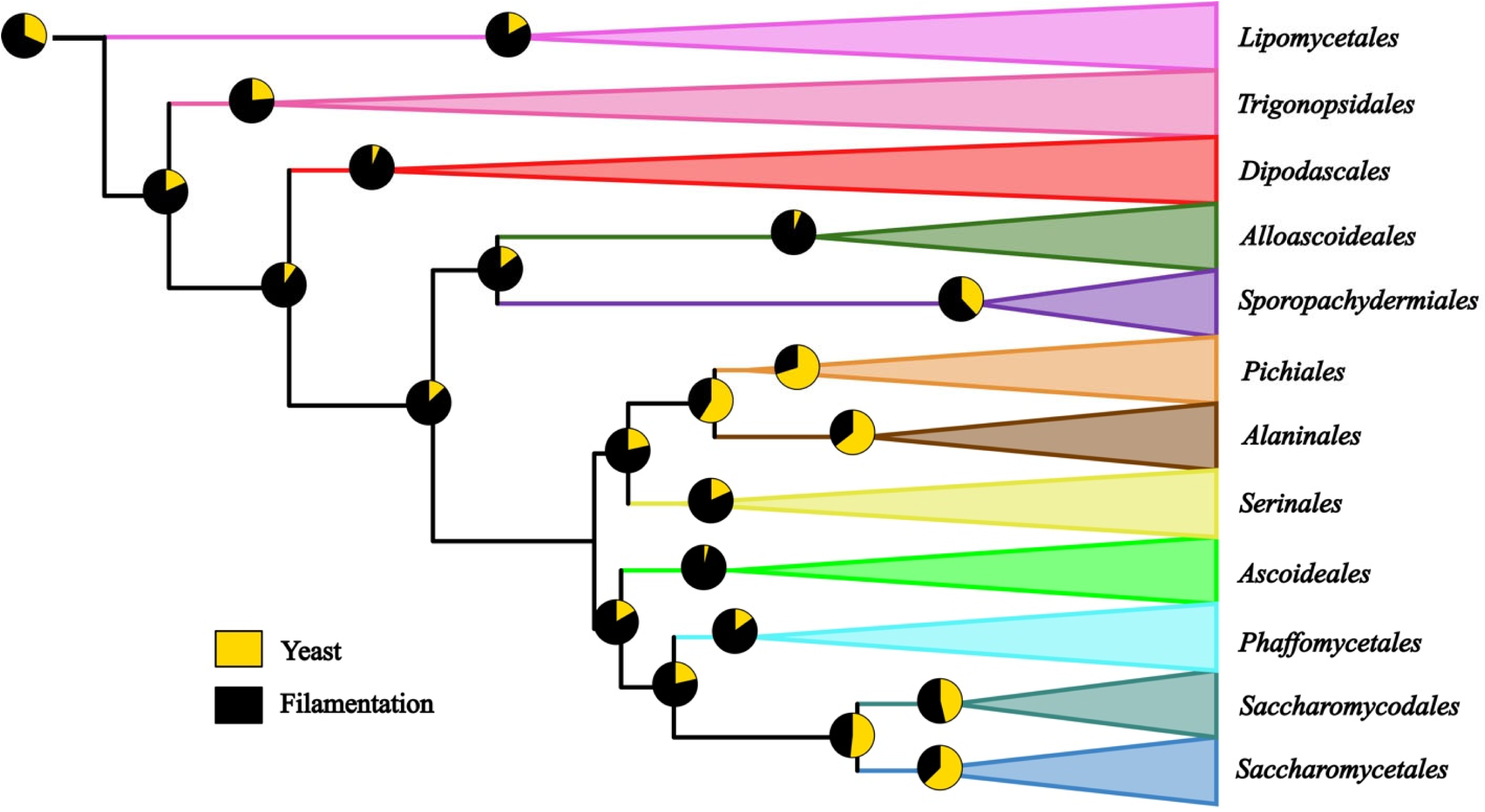
The last common ancestors of most of the 12 taxonomic orders of *Saccharomycotina* yeasts likely had the ability for filamentous growth. The legend on the bottom left includes yeast only (yellow), and filamentation (black) which includes the following filamentation traits: yeast and pseudohyphae and true hyphae, yeast and true hyphae, pseudohyphae, pseudohyphae and true hyphae, and true hyphae. Each taxonomic order is collapsed as a triangle; only the ancestral state likelihood pie charts for the last common ancestors of the 12 orders and their ancestors are shown, and at the edges of collapsed orders are each of the order names. Each different color represents a different taxonomic order.

It is well established that model species across fungi utilize both yeast and filamentous lifestyles to adapt to their surrounding environment (Berman, 2006; Karasz et al., 2022; Naranjo-Ortiz & Gabaldón, 2019). Filamentous growth across *Saccharomycotina* yeasts was predicted with moderate accuracy from genomic features (71% accuracy) and metabolic features (67% accuracy) using machine learning. Genomic features associated with filamentation were related to fatty acid metabolism and cell polarity, while metabolic features included carbon (e.g., α-glucosides) and nitrogen sources (Figure 3). Growth on maltose, an α-glucoside that was the second most important metabolic feature, has been shown to result in elongated cell growth and induced filamentous MAPK signaling relative to glucose (Vandermeulen & Cullen, 2023).

What are the evolutionary and ecological dimensions of this extensive variation in the cellular morphologies of *Saccharomycotina* species? It is well established that ecologically relevant compounds can trigger filamentation in model yeasts, such as *S. cerevisiae* (Vandermeulen & Cullen, 2023). Experimental evolution of unicellular *S. cerevisiae* from clump-forming ancestors showed that clump-formers tend to be more resistant to a variety of stressors than their unicellular counterparts at the cost of slower growth (Kuzdzal-Fick et al., 2019). While it is tempting to assume that the different morphologies likely offer selective advantages in particular environments, recent laboratory evolution experiments by Farkas and coworkers showed that new cellular morphologies can emerge as a byproduct of compensatory evolution (Farkas et al., 2022).

In summary, examination of filamentation across *Saccharomycotina* showed that it is widespread and that true hyphal, pseudohyphal, and yeast morphologies are common. The abundance of observed variation raises several interesting questions, including: Are there specific environmental and/or nutritional conditions that favor filamentous vs. yeast growth? What are the molecular mechanisms that govern polymorphism within individual species? What evolutionary changes gave rise to the observed variation in morphologies across the subphylum? The availability of genomic, metabolic, and ecological data for nearly every species of *Saccharomycotina* (M. Harrison et al., 2024; Opulente et al., 2024) coupled with subphylum-level systems and assays for dissecting gene and pathway functions (e.g., broadly active state-of-the-art genome-editing tools (Kuang et al., 2018)) and genome-enabled multi-omics (Aranguiz et al., 2024; Horianopoulos et al., 2025) make the subphylum a superb system for understanding the relationship between two hallmark fungal morphologies.

## Methods

### Data collection from theyeasts.org

Kurtzman et al. (2011) and its online resource theyeasts.org contain information on the growth profiles of the type strains of all known yeast species, including most *Saccharomycotina* yeast species, as well as descriptions of their asexual, sexual, and filamentous morphologies. The Dalmau plate culture, which uses cornmeal agar and incubation temperatures between 20° C and 25° C to create a microaerobic environment that induces stress and the formation of filaments, was used for most strains to induce filamentous growth; if the yeast grew filaments of pseudohyphae and true hyphae, their presence and further details of growth type were included.

To induce yeast growth, either glucose-yeast extract-peptone agar or YM broth or agar (4% malt extract / 0.5% yeast extract agar). For each *Saccharomycotina* yeast species, filamentation types were input into a morphology trait matrix (MTM) in which filamentous traits were formatted as binary absence / presence traits. Some species were described in theyeasts.org to generate both pseudohyphae and true hyphae and were considered polymorphic. Some species were not included in theyeasts.org; thus, the total number of strains was 1,021 from 979 species of *Saccharomycotina* yeasts.

It is important to note that the morphological traits, i.e., the presence of pseudohyphae, true hyphae and yeasts, used for this study were observed in using growth in Dalmau plate cultures and YM growth media. Dalmau plate cultures have been shown to induce hyphal growth, and YM media are commonly used for yeast culture and growth, but there are other media that were not considered in this study. Furthermore, the life cycles are not known for all 1,154 Saccharomycotina. Therefore, the pattern of trait presence / absence in some strains may differ when other growth media are used.

### Variation of filamentation across *Saccharomycotina* yeasts

Categorical values from the MTM were used as data input for iTOL software version 7 (Letunic & Bork, 2024) to generate a visual map of filamentation types across the entire sub-phylum of *Saccharomycotina* yeasts. The frequency of filamentation was quantified as frequencies per number of species in each specified order. For example, *Lipomycetales* order has data from 36 yeasts; of those, three are not included in the database (8.33%), 29 do not form filaments (87.88%), one can produce pseudohyphae (3.03%), three can produce true hyphae (9.09%), and none are polymorphic for pseudohyphae and true hyphae (0%). Histograms with the counts and frequencies of the different types of morphologies were generated in Microsoft Excel 2024.

### Phylogenetic analysis of filamentation types

For phylogenetic signal and ancestral state reconstruction, we used Picante R-package version 1.8.2 (Kembel et al., 2010) function match.phylo.data() to match each trait data file (either binary data for phylogenetic signal, or categorical data for ancestral state reconstruction) to the phylogeny. Phylogenetic signal was calculated for various binary filamentation traits (i.e., presence / absence of filaments, presence / absence of pseudohyphae, presence / absence of true hyphae, presence / absence of pseudohyphae and true hyphae) using the D-statistic (Fritz & Purvis, 2010). Caper software version 1.0.3 (Orme et al., 2023) function phylo.d() was used to calculate the D-statistic value and associated p-values (under Brownian motion model vs. under no, i.e., random, phylogenetic structure) for each filamentation type. To control the random permutations, a fixed seed was used for result reproducibility.

### Ancestral state reconstruction of filamentation

To estimate the evolutionary history of all *Saccharomycotina* yeast internal nodes, ancestral state reconstruction was used. The transition rates for the transitions between each filament trait state was estimated with Phytools version 2.4-4 (Revell, 2024) function fitpolyMk, which estimates discrete character evolution using a Markov model (Lewis, 2001).

Some species are described to be polymorphic and generate both pseudohyphae and true hyphae; thus, the unordered transitions between each state (YEAST, PSEUDOHYPHAE, and TRUE HYPHAE) were needed to include all hidden intermediate states (YEAST + PSEUDOHYPHAE, YEAST + TRUE HYPHAE, YEAST + PSEUDOHYPHAE + TRUE HYPHAE, and PSEUDOHYPHAE + TRUE HYPHAE). Transition rates were estimated under four transition rate models: ER (equal rates), ARD (all rates different), SYM (symmetric backward and forward rates), and transient (polymorphisms are gained at different rates than polymorphisms are lost). Model selection was performed using the anova() function (Chambers and Hastie 1992), and the model with the lowest Akaike Information Criterion (AIC) was chosen for downstream analysis.

The best-fitting transition rate model was implemented in a hidden Markov model using the corHMM package version 2.8 (Beaulieu et al., 2013). This model allows for rate variation and estimates the likelihood of each trait state being present at all ancestral nodes of the phylogeny. The resulting likelihood values were used to calculate the probability of each specific trait state at each ancestral node. These probabilities were then visualized as pie charts mapped across the phylogeny.

### Genomic data matrix

InterProScan gene functional annotations were generated by the Y1000+ Project (Opulente et al., 2024). A data matrix was built with counts of the unique InterPro ID number for each *Saccharomycotina* strain genome, in which rows were strains and columns contained the count of each InterPro ID (N = 12,242) present in one or more of the 1,154 yeast genomes. The number of every InterPro ID for each genome was calculated with a python script and input into corresponding cells in the data matrix (Blum et al., 2025).

### Metabolic data matrix

The metabolic data matrix comprised 122 traits measured across 893 yeast strains of the total 1,154 representing 885 species within the subphylum (M.-C. Harrison et al., 2024; Opulente et al., 2018). The matrix contained traits across a range of growth capacities on carbon and nitrogen sources (e.g. galactose, raffinose, urea), as well as environmental conditions (e.g. growth at different temperatures and salt concentrations). Traits that were less frequently studied had higher levels of missing data compared to more commonly studied traits; the average percentage of missing data was 37.5% (40,906 of 108,946 total entries).

### Environmental data matrix

The isolation environments were collected for 94% of all yeasts (1,088 of 1,154 strains) sourced from strain databases, species descriptions, or from *The Yeasts: A Taxonomic Study* (Kurtzman et al., 2011; Opulente et al., 2024). To include all unique isolation environment descriptions, the environmental data were encoded into a hierarchical binary trait matrix using a controlled vocabulary (M. Harrison et al., 2024; Opulente et al., 2024). The ecology ontology contains six broad categories of isolation environments including animal, plant, environmental, fungal, industrial products, and victuals (food or drink), with specific controlled vocabulary used to annotate each strain.

### Machine learning methods to predict filamentation across *Saccharomycotina* yeasts

We used a random forest algorithm to determine if the presence and / or absence of filamentation could be predicted from genomic, metabolic, or environmental features. Independent random forest models were trained on the different datasets to assess classification accuracy and identify the top predictive features. While the function of the model is classification, and the random forest algorithm that we use is a classifier, we refer to the outcomes as “predictions” for ease of interpretation.

A machine learning model was trained using the XGBoost version 1.7.3 (Orme et al., 2023.) random forest classifier (XGBRFClassifier()) with the parameters max_depth=12 and n_estimators=100; all other parameters were in their default settings. To avoid overfitting and preserve the highest accuracy, the max_depth parameter determines how complex the random forest will be by setting the depth of each decision tree. The number of decision trees in the model was indicated with the n_estimators parameter. Evaluation of impact of increasing each parameter showed that higher max_depth and more decision trees per random forest model further improved accuracy.

The sklearn.model_selection version 1.2.1 package functions RepeatedStratifiedKFold and cross_val_score were used to train 90% of the data and cross validation of the remaining 10% for the random forest model (Pedregosa et al., 2011). Cross validation is an accuracy assessment method that involves 10 trials, each time holding back a random 10% of the training data for model testing. The Sci-Kit Learn function cross_val_predict() was used independently to generate the confusion matrices; matrices show the amount of yeasts predicted correctly to either be capable of generating filamentous growth or only yeast growth (true positives and true negatives, respectively) and predicted incorrectly (false positives, predicted to be filamentous but are in reality do not generate filamentous growth (exist only as yeast); and false negatives, predicted to only have yeast growth but are in reality also generate filaments). The cross_val_predict() function also performs 10-fold cross validation, tracking which species are true / false positives and those that are true / false negatives for each of the 10 trials and compiling the final results into a confusion matrix. The XGBRFClassifier function was used to generate the top predictive features using Gini importance, which is based on node impurity (the amount of variance in filamentous growth with strains that either generate filaments or not). We recorded and saved all metrics, including total balanced accuracy (in which 50% accuracy is interpreted as randomly guessing).

Receiver Operating Characteristic (ROC) curves, which plot the true positive rate against the false positive rate, were generated for each prediction analysis to visualize the accuracy of the algorithm in predicting filamentous growth. An area under the curve (AUC) greater than 0.5 indicates classification better than random. To capture the error that could exist in the whole dataset, non-down-sampled datasets were used for the ROC analysis.

## Acknowledgements

We thank members of the Rokas lab and the Y1000+ Project (http://y1000plus.org) for helpful discussions. This project was supported by the National Science Foundation under Grants Nos. DEB-2110403 (to C.T.H.) and DEB-2110404 (to A.R.); in part by the Great Lakes Bioenergy Research Center, U.S. Department of Energy, Office of Science, Biological and Environmental Research Program under Award Number DE-SC0018409 (of which C.T.H. is a co-investigator); and the National Institute of Food and Agriculture, United States Department of Agriculture, Hatch project 7005101 (to C.T.H.). C.T.H. is a Vilas Faculty Mid-Career Investigator. Research in A.R.’s lab is also supported by the National Institutes of Health/National Institute of Allergy and Infectious Diseases (R01 AI153356).

## Author contributions

C.M.C and A.R. conceptualized this project. C.M.C. and T.D. developed the methodology for ancestral state reconstruction, and M.C.H. developed the methodology for machine learning. C.M.C. curated and analyzed the data, as well as prepared the figures under the supervision and guidance of A.R. C.M.C wrote the manuscript, and M.C.H., T.D., M.G., A.R., and C.T.H. reviewed edited and approved the manuscript. A.R. and C.T.H. acquired funding for this project.

## Conflicts of Interest

A.R. is a scientific consultant of LifeMine Therapeutics, Inc. All other authors declare no competing interests.

## References

Aranguiz, K., Horianopoulos, L. C., Elkin, L., Abá, K. S., Wrobel, R. L., Shiu, S.-H., Rokas, A., & Hittinger, C. T. (2024). Machine learning reveals genes impacting oxidative stress resistance across yeasts. Microbiology. 10.1101/2024.08.14.607963

Basso, V., d’Enfert, C., Znaidi, S., & Bachellier-Bassi, S. (2019). From genes to networks: The regulatory circuitry controlling *Candida albicans* morphogenesis. Current Topics in Microbiology and Immunology, 422, 61–99. 10.1007/82_2018_144

Beaulieu, J. M., O’Meara, B. C., & Donoghue, M. J. (2013). Identifying Hidden Rate Changes in the Evolution of a Binary Morphological Character: The Evolution of Plant Habit in Campanulid Angiosperms. Systematic Biology, 62(5), 725–737. 10.1093/sysbio/syt034

Berman, J. (2006). Morphogenesis and cell cycle progression in Candida albicans. Current Opinion in Microbiology, 9(6), 595–601. 10.1016/j.mib.2006.10.007

Bi, E., & Park, H.-O. (2012). Cell Polarization and Cytokinesis in Budding Yeast. Genetics, 191(2), 347–387. 10.1534/genetics.111.132886

Blum, M., Andreeva, A., Florentino, L. C., Chuguransky, S. R., Grego, T., Hobbs, E., Pinto, B. L., Orr, A., Paysan-Lafosse, T., Ponamareva, I., Salazar, G. A., Bordin, N., Bork, P., Bridge, A., Colwell, L., Gough, J., Haft, D. H., Letunic, I., Llinares-López, F.,… Bateman, A. (2025). InterPro: The protein sequence classification resource in 2025. Nucleic Acids Research, 53(D1), D444–D456. 10.1093/nar/gkae1082

Chavez, C. M., Groenewald, M., Hulfachor, A. B., Kpurubu, G., Huerta, R., Hittinger, C. T., & Rokas, A. (2024). The cell morphological diversity of *Saccharomycotina* yeasts. FEMS Yeast Research, 24, foad055. 10.1093/femsyr/foad055

Chen, T., & Guestrin, C. (2016). XGBoost: A Scalable Tree Boosting System. Proceedings of the 22nd ACM SIGKDD International Conference on Knowledge Discovery and Data Mining, 785–794. 10.1145/2939672.2939785

Chiou, J., Balasubramanian, M. K., & Lew, D. J. (2017). Cell Polarity in Yeast. Annual Review of Cell and Developmental Biology, 33(1), 77–101. 10.1146/annurev-cellbio-100616-060856

Chow, E. W. L., Pang, L. M., & Wang, Y. (2021). From Jekyll to Hyde: The Yeast–Hyphal Transition of Candida albicans. Pathogens, 10(7), 859. 10.3390/pathogens10070859

Crandall, J. G., Zhou, X., Rokas, A., & Hittinger, C. T. (2024). Specialization Restricts the Evolutionary Paths Available to Yeast Sugar Transporters. Molecular biology and evolution, 41(11), msae228. 10.1093/molbev/msae228

Cullen, P. J., & Sprague, G. F. (2000). Glucose depletion causes haploid invasive growth in yeast. Proceedings of the National Academy of Sciences, 97(25), 13619–13624. 10.1073/pnas.240345197

Cullen, P. J., & Sprague, G. F. (2012). The Regulation of Filamentous Growth in Yeast. Genetics, 190(1), 23–49. 10.1534/genetics.111.127456

David, K. T., Schraiber, J. G., Crandall, J. G., Labella, A. L., Opulente, D. A., Harrison, M.-C., Wolters, J. F., Zhou, X., Shen, X.-X., Groenewald, M., Hittinger, C. T., Pennell, M., & Rokas, A. (2025). Convergent expansions of keystone gene families drive metabolic innovation in Saccharomycotina yeasts. Proceedings of the National Academy of Sciences, 122(23), e2500165122. 10.1073/pnas.2500165122

Diepeveen, E. T., De La Cruz, L. I., & Laan, L. (2017). Evolutionary dynamics in the fungal polarization network, a mechanistic perspective. Biophysical Reviews, 9(4), 375–387. 10.1007/s12551-017-0286-2

Du, H., Bing, J., Hu, T., Ennis, C. L., Nobile, C. J., & Huang, G. (2020). Candida auris: Epidemiology, biology, antifungal resistance, and virulence. PLOS Pathogens, 16(10), e1008921. 10.1371/journal.ppat.1008921

Farkas, Z., Kovács, K., Sarkadi, Z., Kalapis, D., Fekete, G., Birtyik, F., Ayaydin, F., Molnár, C., Horváth, P., Pál, C., & Papp, B. (2022). Gene loss and compensatory evolution promotes the emergence of morphological novelties in budding yeast. Nature Ecology & Evolution, 6(6), 763–773. 10.1038/s41559-022-01730-1

Fritz, S. A., & Purvis, A. (2010). Selectivity in Mammalian Extinction Risk and Threat Types: A New Measure of Phylogenetic Signal Strength in Binary Traits. Conservation Biology, 24(4), 1042–1051. 10.1111/j.1523-1739.2010.01455.x

Groenewald, M., Hittinger, C. T., Bensch, K., Opulente, D. A., Shen, X.-X., Li, Y., Liu, C., LaBella, A. L., Zhou, X., Limtong, S., Jindamorakot, S., Gonçalves, P., Robert, V., Wolfe, K. H., Rosa, C. A., Boekhout, T., Ĉadež, N., Péter, G., Sampaio, J. P.,… Rokas, A. (2023). A genome-informed higher rank classification of the biotechnologically important fungal subphylum *Saccharomycotina*. Studies in Mycology, 105(1), 1–22. 10.3114/sim.2023.105.01

Harris, S. D. (2011). Hyphal morphogenesis: An evolutionary perspective. Fungal Biology, 115(6), 475–484. 10.1016/j.funbio.2011.02.002

Harrison, M., Opulente, D. A., Wolters, J. F., Shen, X., Zhou, X., Groenewald, M., Hittinger, C. T., Rokas, A., & LaBella, A. L. (2024). Exploring Saccharomycotina Yeast Ecology Through an Ecological Ontology Framework. Yeast, 41(10), 615–628. 10.1002/yea.3981

Harrison, M.-C., Ubbelohde, E. J., LaBella, A. L., Opulente, D. A., Wolters, J. F., Zhou, X., Shen, X.-X., Groenewald, M., Hittinger, C. T., & Rokas, A. (2024). Machine learning enables identification of an alternative yeast galactose utilization pathway. Proceedings of the National Academy of Sciences, 121(18), e2315314121. 10.1073/pnas.2315314121

Horianopoulos, L. C., Rokas, A., & Hittinger, C. T. (2025). Convergent evolution of aerobic fermentation through divergent mechanisms acting on key shared glycolytic genes. bioRxiv : the preprint server for biology, 2025.10.02.679963. 10.1101/2025.10.02.679963

Il’chenko, A. P., Ogorelyshev, D. I., Shyshkanova, N. V., Sokolov, A. P., Finogenova, T. V., & Kondrashova, M. N. (2005). The Effect of Succinate on Respiration, Transamination, and Pyruvate Formation in Cells of the Yeast Dipodascus magnusii. Microbiology, 74(5), 527–532. 10.1007/s11021-005-0099-3

Karasz, D. C., Weaver, A. I., Buckley, D. H., & Wilhelm, R. C. (2022). Conditional filamentation as an adaptive trait of bacteria and its ecological significance in soils. Environmental Microbiology, 24(10), 4966–4966. 10.1111/1462-2920.16228

Kembel, S. W., Cowan, P. D., Helmus, M. R., Cornwell, W. K., Morlon, H., Ackerly, D. D., Blomberg, S. P., & Webb, C. O. (2010). Picante: R tools for integrating phylogenies and ecology. Bioinformatics, 26(11), 1463–1464. 10.1093/bioinformatics/btq166

Kiss, E., Hegedüs, B., Virágh, M., Varga, T., Merényi, Z., Kószó, T., Bálint, B., Prasanna, A. N., Krizsán, K., Kocsubé, S., Riquelme, M., Takeshita, N., & Nagy, L. G. (2019). Comparative genomics reveals the origin of fungal hyphae and multicellularity. Nature Communications, 10(1), 4080. 10.1038/s41467-019-12085-w

Kuang, M. C., Kominek, J., Alexander, W. G., Cheng, J.-F., Wrobel, R. L., & Hittinger, C. T. (2018). Repeated Cis-Regulatory Tuning of a Metabolic Bottleneck Gene during Evolution. Molecular Biology and Evolution, 35(8), 1968–1981. 10.1093/molbev/msy102

Kurtzman, C. P., Fell, J. W., & Boekhout, T. (2011). The Yeasts, a Taxonomic Study. (5th ed.). Elsevier Science.

Kuzdzal-Fick, J. J., Chen, L., & Balázsi, G. (2019). Disadvantages and benefits of evolved unicellularity versus multicellularity in budding yeast. Ecology and Evolution, 9(15), 8509–8523. 10.1002/ece3.5322

Lambrechts, M. G., Bauer, F. F., Marmurt, J., & Pretorius, I. S. (1996). Mucl, a mucin-like protein that is regulated by MsslO, is critical for pseudohyphal differentiation in yeast. Proc. Natl. Acad. Sci. USA.

Laxman, S., & Tu, B. P. (2011). Multiple TORC1-Associated Proteins Regulate Nitrogen Starvation-Dependent Cellular Differentiation in Saccharomyces cerevisiae. PLoS ONE, 6(10), e26081. 10.1371/journal.pone.0026081

Letunic, I., & Bork, P. (2024). Interactive Tree of Life (iTOL) v6: Recent updates to the phylogenetic tree display and annotation tool. Nucleic Acids Research, 52(W1), W78–W82. 10.1093/nar/gkae268

Lewis, P. O. (2001). A Likelihood Approach to Estimating Phylogeny from Discrete Morphological Character Data. Systematic Biology, 50(6), 913–925. 10.1080/106351501753462876

Matheos, D., Metodiev, M., Muller, E., Stone, D., & Rose, M. D. (2004). Pheromone-induced polarization is dependent on the Fus3p MAPK acting through the formin Bni1p. The Journal of Cell Biology, 165(1), 99–109. 10.1083/jcb.200309089

Mourer, T., El Ghalid, M., d’Enfert, C., & Bachellier-Bassi, S. (2021). Involvement of amyloid proteins in the formation of biofilms in the pathogenic yeast Candida albicans. Research in Microbiology, 172(3), 103813. 10.1016/j.resmic.2021.103813

Mukaremera, L., Lee, K. K., Mora-Montes, H. M., & Gow, N. A. R. (2017). Candida albicans Yeast, Pseudohyphal, and Hyphal Morphogenesis Differentially Affects Immune Recognition. Frontiers in Immunology, 8, 629. 10.3389/fimmu.2017.00629

Naranjo-Ortiz, M. A., & Gabaldón, T. (2019). Fungal evolution: Diversity, taxonomy and phylogeny of the Fungi. Biological Reviews, 94(6), 2101–2137. 10.1111/brv.12550

Opulente, D. A., LaBella, A. L., Harrison, M.-C., Wolters, J. F., Liu, C., Li, Y., Kominek, J., Steenwyk, J. L., Stoneman, H. R., VanDenAvond, J., Miller, C. R., Langdon, Q. K., Silva, M., Gonçalves, C., Ubbelohde, E. J., Li, Y., Buh, K. V., Jarzyna, M., Haase, M. A. B.,… Hittinger, C. T. (2024). Genomic factors shape carbon and nitrogen metabolic niche breadth across Saccharomycotina yeasts. Science, 384(6694), eadj4503. 10.1126/science.adj4503

Opulente, D. A., Rollinson, E. J., Bernick-Roehr, C., Hulfachor, A. B., Rokas, A., Kurtzman, C. P., & Hittinger, C. T. (2018). Factors driving metabolic diversity in the budding yeast subphylum. BMC Biology, 16(1), 26. 10.1186/s12915-018-0498-3

Orme, D., Freckleton, R., Thomas, G., Petzoldt, T., Fritz, S., Isaac, N., & Pearse, W. (2023). caper: Comparative Analyses of Phylogenetics and Evolution in R (R package version 1.0.3)

Pedregosa, F., Varoquaux, G., Gramfort, A., Michel, V., Thirion, B., Grisel, O., Blondel, M., Prettenhofer, P., Weiss, R., Dubourg, V., Vanderplas, J., Passos, A., Cournapeau, D., Brucher, M., Perrot, M., & Duchesnay, É. (2011). Scikit-learn: Machine learning in Python. Journal of Machine Learning Research, 12, 2825–2830.

Pérez-Campo, F. M., & Domínguez, A. (2001). Factors Affecting the Morphogenetic Switch in Yarrowia lipolytica. Current Microbiology, 43(6), 429–433. 10.1007/s002840010333

Pérez-Izquierdo, L., Rincón, A., Lindahl, B. D., & Buée, M. (2021). Fungal community of forest soil: Diversity, functions, and services. In Forest Microbiology (pp. 231–255). Elsevier. 10.1016/B978-0-12-822542-4.00022-X

Powers-Fletcher, M. V., Kendall, B. A., Griffin, A. T., & Hanson, K. E. (2016). Filamentous Fungi. Microbiology Spectrum, 4(3), 4.3.23. 10.1128/microbiolspec.DMIH2-0002-2015

Revell, L. J. (2024). phytools 2.0: An updated R ecosystem for phylogenetic comparative methods (and other things). PeerJ, 12, e16505. 10.7717/peerj.16505

Sudbery, P. E. (2011). Growth of Candida albicans hyphae. Nature Reviews Microbiology, 9(10), 737–748. 10.1038/nrmicro2636

Váchová, L., & Palková, Z. (2018). How structured yeast multicellular communities live, age and die? FEMS Yeast Research, 18(4). 10.1093/femsyr/foy033

Van De Velde, S., & Thevelein, J. M. (2008). Cyclic AMP-Protein Kinase A and Snf1 Signaling Mechanisms Underlie the Superior Potency of Sucrose for Induction of Filamentation in *Saccharomyces cerevisiae*. Eukaryotic Cell, 7(2), 286–293. 10.1128/EC.00276-07

Van Mulders, S. E., Christianen, E., Saerens, S. M. G., Daenen, L., Verbelen, P. J., Willaert, R., Verstrepen, K. J., & Delvaux, F. R. (2009). Phenotypic diversity of Flo protein family-mediated adhesion in *Saccharomyces cerevisiae*. FEMS Yeast Research, 9(2), 178–190. 10.1111/j.1567-1364.2008.00462.x

Vandermeulen, M. D., & Cullen, P. J. (2023). Ecological inducers of the yeast filamentous growth pathway reveal environment-dependent roles for pathway components. mSphere, 8(5), e00284–23. 10.1128/msphere.00284-23

Verstrepen, K. J., & Klis, F. M. (2006). Flocculation, adhesion and biofilm formation in yeasts. Molecular Microbiology, 60(1), 5–15. 10.1111/j.1365-2958.2006.05072.x

Wu, B., Hao, W., & Cox, M. P. (2022). Reconstruction of gene innovation associated with major evolutionary transitions in the kingdom Fungi. BMC Biology, 20(1), 144. 10.1186/s12915-022-01346-8

